# Hyphal growth determines spatial organization and coexistence in a pathogenic polymicrobial community within alveoli-like geometries

**DOI:** 10.1101/2025.03.22.644766

**Authors:** L Mancini, L Saliekh, R Claydon, J Kotar, E Benyei, C A Munro, T N Shendruk, A Brown, M Welch, P Cicuta

## Abstract

The bodies of macroorganisms host microbes living in multi-species communities. Sequencing approaches have revealed that different organs host different microbiota and tend to be infected by different pathogens, drawing correlations between environmental parameters at the organ level and microbial composition. However, less is known about the microscale dimension of microbial ecology, particularly during infection. In this study, we focus on the role of microscale spatial structure, studying its influence on the ecology of a polymicrobial infection of *P. aeruginosa, S. aureus* and *C. albicans*. Although these pathogens are commonly found together in the lungs of chronically ill patients, it is unclear whether they coexist or compete and segregate in different niches. We find that, while *P. aeruginosa* quickly outcompetes *C. albicans* and *S. aureus* on large surfaces, robust spatial organization and coexistence emerges in microfluidic microchambers that mimic the spatial characteristics of alveoli. In these microenvironments, slowly growing *C. albicans* is able to leverage fast eccentric hyphal growth to conquer boundary spaces, where it establishes itself excluding the other pathogens. We show that the emerging spatial patterning is robust to changes in the virulence of the community, enabling coexistence and potentially determining infection severity and outcomes. Our findings reveal a previously unrecognized role of mechanical forces in shaping infection dynamics, suggesting that microenvironmental structure is a critical determinant of pathogen coexistence, virulence, and treatment outcomes. Because adaptations, such as changes in morphology, are widespread among microbes, these results are generalizable to other ecologies and environments.

## INTRODUCTION

“Omic” studies have revealed that microbes seldom live in isolation, both in the healthy (1; 2; 3) and diseased body (4; 5; 6). Instead, many different species come together to consume substrates giving rise to polymicrobial communities and rich ecologies (7; 8; 9). Interactions with different structural characteristics of their environments produce spatial patterning that in “macroorganisms” we commonly call biogeography (10). This patterning on both the macro and microscale eventually determines the phenotypic outcome of the polymicrobial community, either in terms of virulence (11; 12; 13; 14; 15; 16), survival to treatment (17; 18; 19; 20), or substrate transformation (21; 22; 23; 24).

The biogeography of microbes is influenced by a number of factors. The effects of anatomy, host immune system, physiology and abiotic conditions (pH, oxygenation) are the most likely to be detected by “omics”, because they vary on larger scales and are therefore better known (25). These factors determine, for example, the composition of the organ-specific microbiota (26; 27). Biotic factors and ecological interactions between microbes tend to act on the microscale (25) and have been studied mainly through mathematical modeling (28; 29) or in synthetic communities grown in the lab (30; 31; 32; 33; 34). In addition to revealing a wealth of new ecology, these “synthetic” communities have the merit of enabling live direct observations.

Microscopy compensates for several of the shortcomings of “omic” approaches, such as the limited spatial resolution and their insensitivity to morphology and behavior. In addition, while sequencing approaches can robustly draw associations between the presence of certain microbes and pathological conditions, they provide limited information to establish causal links and mechanisms. Direct observation unlocks access to the microscale, where microbe-microbe and host-microbe physical and molecular interactions occur. Despite progress in biomedical imaging, the observation of these microbial consortia *in vivo* in humans remains a challenging task. Some indications on the shapes and forms of these communities are provided by *ex vivo* observations. These images have been fundamental to understanding that microbes can indeed have complex spatial organizations at the microscale - an “urban biogeography” - that can have significant consequences on their pathogenicity and biophysical and metabolic properties (35; 22). Understanding how specific “urban biogeographies” come about and mapping different organizations to their emergent properties will open up new intervention strategies aimed at preventing and treating pathogenic scenarios, including infection.

To achieve this goal, approaches that encompass the host, polymicrobial communities, live microscopy, and controlled environments are needed. Currently, two model scenarios satisfy these requirements in different measures and their choice depends on the research question: animal models and infection-on-chip. Animal models offer the most realistic conditions as they respond to infection with a complete immune system and therefore are well suited to study the complex interaction between microbes and host response (36; 37; 38). However, imaging infection in these systems is challenging (39). Even in transparent hosts, such as zebrafish larvae, the spatial resolution for live microscopy is limited to the organ level (40; 41; 42; 43; 44; 45; 46). Organoids and infection-on-chip systems that encompass host tissues strike a compromise, improving imaging to the detriment of the immune response, which tends to be only partial (47; 48; 49; 50). In both cases, complexity remains high and isolating the mechanistic contribution of specific parameters can be challenging.

In this work, we focus on the role of spatial structure in the lung on shaping the microscale ecology of a respiratory pathogenic polymicrobial community. A large body of work (51; 52; 53) has demonstrated that microbial phenotype and evolution are conditioned by the mechanical properties of the substrate, both passively (54) and through active sensing (55; 56). Contact with surfaces is commonly associated with the formation of mono- and polymicrobial biofilms (57; 58), which are studied almost exclusively on surfaces that do not impose limitations in the surface plane (59). In these settings, spatial organization within biofilms has been studied as an emergent property of microbe-microbe interactions, with surfaces acting as a point of anchorage that limits growth and movement in the direction perpendicular to the surface (13; 60; 61; 62). The microenvironments within host which microbes inhabit are often more complex than simple planar surfaces, and thus these models neglect important mechanical contributions (63; 54). Because of these limitations, questions regarding the impact of the mechanical and spatial characteristics of the host microenvironment on the ecology of polymicrobial communities remain open. For example, it is unclear why the trio of microbes *P. aeruginosa, S. aureus* and *C. albicans*, that are difficult to co-culture in the lab, (64) are consistently found together in the sputum of patients, particularly those affected by chronic diseases such as cystic fibrosis (65; 66; 67; 68; 69). Do different species in the lung coexist or are they segregated in different regions of the organ?

To investigate the impact of spatial structure on microbial ecology, we compare the spatial organization of polymicrobial communities of *P. aeruginosa, S. aureus* and *C. albicans* in two scenarios: unconfined surfaces and alveoli mimics. With the latter, we expand the definition of an infection-on-chip, producing microfluidics that do not contain host cells but that reproduce the biochemical properties of the infection site and its key spatial features. This allows us to specifically control and isolate the impact of confinement on ecology, bypassing confounding factors that emerge from microbe-host interactions or from the requirements of host cell culturing. We find that, although *P. aeruginosa* quickly outcompetes the other species on an unlimited surface, alveoli-like confinement profoundly alters ecology, allowing persistent polymicrobial cohabitation. In the alveoli mimics, we observe the robust emergence of a spatial organization that enables co-existence. Despite growing slower than the other species, *C. albicans* is able to conquer the edges and the closed end of the microcompartments. We demonstrate that this is enabled by a transition to hyphal morphology that inherently focuses directional eccentric growth. Upon reaching a barrier, *C. albicans* tends to bend and reorient, continuing growth along the edge and displacing other microbes, as reproduced in our purely biomechanical agent-based simulations. Finally, using a clinically relevant *mexT* mutant of *P. aeruginosa*, we show that the spatial organization observed in the community is robust to changes in virulence of its members. Taken together, our results reveal a previously unrecognized role of mechanical forces in shaping infection dynamics and suggest that the microenvironmental structure is a critical determinant of pathogen coexistence, virulence, and treatment outcomes.

## RESULTS

### P. aeruginosa quickly outcompetes S. aureus and C. albicans on a surface

We consider a lung-relevant pathogenic polymicrobial community of *P. aeruginosa* (PA), *S. aureus* (SA) and *C. albicans* (CA) on the simplest and best studied (23) example of a structured environment: a soft agarose surface. The species in our community expressed different fluorescent proteins that allowed their tracking in epifluorescence microscopy: CFP for PA, GFP for SA and dTomato for CA. To recapitulate aspects of lung biochemistry, plates are composed of artificial sputum medium (ASM) (64). Using a micropipette, we seed small aliquots of cells from the three species mixed in different ratios and observe their growth at 37°C after 24 hours. When the inoculum contained the same number of cells of the three species, despite the much lower cell size and starting biomass, *P. aeruginosa* dominated the mixed colony after 24 hours. The colonies nearly doubled their radii over the time interval. While *C. albicans* and *S. aureus* show limited growth and remain confined to their seeding spots, all of the radius gained by the colony over the 24 hours is due to *P. aeruginosa* growth (Fig. 1A). This is consistent with reports indicating that motility is a major driver of growth on surfaces (70; 71). To understand the extent of such advantage, we repeat the experiment at two more PA:SA:CA seeding ratios: 0.1:0.1:1 and 0.01:0.01:1. Reducing the PA titer by 10 times does not significantly affect the colony growth phenotype observed, with PA still dominating (Fig. 1B). The phenotype switches to CA dominance when its initial titer is 100 times higher (Fig. 1). Interestingly, the colony radius also doubles over 24 hours, confirming competitive inhibition between PA, CA and SA, as previously observed (64).

**Figure 1.**
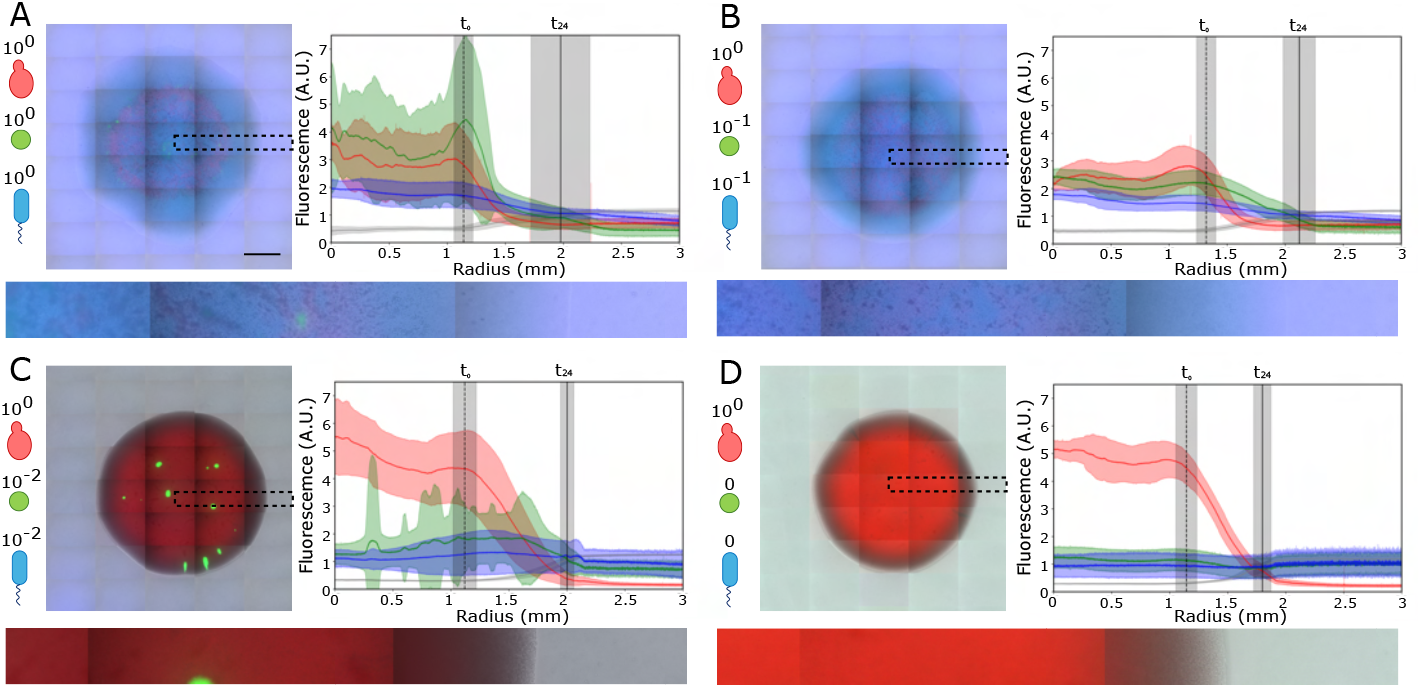
After 24 hours the leading edge of polymicrobial colonies is entirely made up of *P. aeruginosa*. a) PA-SA-CA polymicrobial community seeded at a 1:1:1 ratio on soft agar. (left) example colony after 24 hours growth at 37°C. (bottom) zoom-in of the highlighted area. (right) average intensity profiles for the red, green, blue (fluorescence) and gray (brightfield) channels. Gray reports brightfield intensity and is a proxy for total colony size. To obtain the traces, colonies are split in 360 sections, each 1° large. Radii between the centroid and the edges of the image are drawn and averaged. The shaded areas show the standard deviation of all radii across 5 colonies. The vertical black lines indicate the average radius of the colony at the start and after 24 hours. The shaded area shows the standard deviation. Scale bar = 1 mm. b) PA-SA-CA polymicrobial community seeded at a 0.1:0.1:1 ratio. Averages and standard deviations of 3 colonies. c) b) PA-SA-CA polymicrobial community seeded at a 0.01:0.01:1 ratio. Averages and standard deviations of 3 colonies. d) Growth of a colony of *C. albicans* in isolation. Averages and standard deviations of 3 colonies. Epifluorescence microscopy, stitched FOVs.

### Spatial organization emerges within 10 hours in a lung mimic

Agar surfaces such as the one we used model situations in which space is unlimited in the planar surface and are therefore best suited to study the upper airways. The environment of the lower airways is significantly more structured, with alveolar volumes of the order of millions of cubic micrometers, which, if approximated to spheres, would measure 100 to 250 *µ*m in diameter (72; 73). To model the effects that spatial limits have on the ecology of our pathogenic polymicrobial community, we present a minimal microfluidic model of the distal lung. Our design is inspired by the family machine (74), and encompasses a central channel 1.5 cm long, 1 mm wide, 0.3 mm tall, lined on each side by 100 quasi-2D square alveoli mimics with a side length of 150 *µ*m and a thickness of 8 *µ*m (Fig. 2A and 2B). All measures are modeled on those of the distal lung with the exception of the thickness of the alveolar space, which we chose to limit to 8 *µm* with the intention of observing a single slice of microbial ecology without the need to use confocal microscopy, drastically improving our imaging throughput. The thickness of these alveoli-like spaces allows the stacking of multiple *P. aeruginosa* and *S. aureus* cells, while *C. albicans* cells are limited to one (Fig. 1C, SI Fig. 1). In larger aveoli mimics with side length of 500 *µ*m, a slowdown of growth emerges towards the closed end, indicating the development of nutritional gradients (SI Fig. 2A). Cells are loaded in the alveoli mimics at different species titers (Fig. 2D) and continuously perfused with fresh ASM for up to 20 hours while we perform timelapse microscopy. The observation window is limited to this interval because at later stages robust biofilm growth in the main channel starts influencing cell behavior in the alveoli (SI Fig. 3). Across eight independent experiments (Fig. 2D) and 900 alveoli-mimics, the robust emergence of spatial organization is observed (Fig. 2E and 2G) with a tendency of *C. albicans* to localize towards the closed end and the edges. This displaces *P. aeruginosa* and *S. aureus* and in response they tend to localize towards the center and the entrance of the box. The displacement is most evident in *S. aureus* (Fig. 2H). This spatial organization emerges in less than 10 hours and remains stable throughout the course of the observations (Fig. 2I). As expected, the outcome has a certain dependence on the seeding titer of the species, with high *C. albicans*-to-bacteria ratios leading to dominance by the fungus and vice versa (Fig. 2D). To ensure that fluorescence measurements in the alveoli mimic are a good proxy for the biomass of each species, we carry out control experiments inside species-specific mother machines (SI Fig. 4). Because the two quantities are in good agreement, we utilize the time traces of each fluorescence channel to extract the growth rate of each species in the polymicrobial context within the alveoli mimics. Despite the observed sharing of space and the robust tendency of *C. albicans* to conquer the closed end of the alveoli-mimics, we find significantly different growth rates, with the fungus growing the slowest (SI Fig. 5).

**Figure 2.**
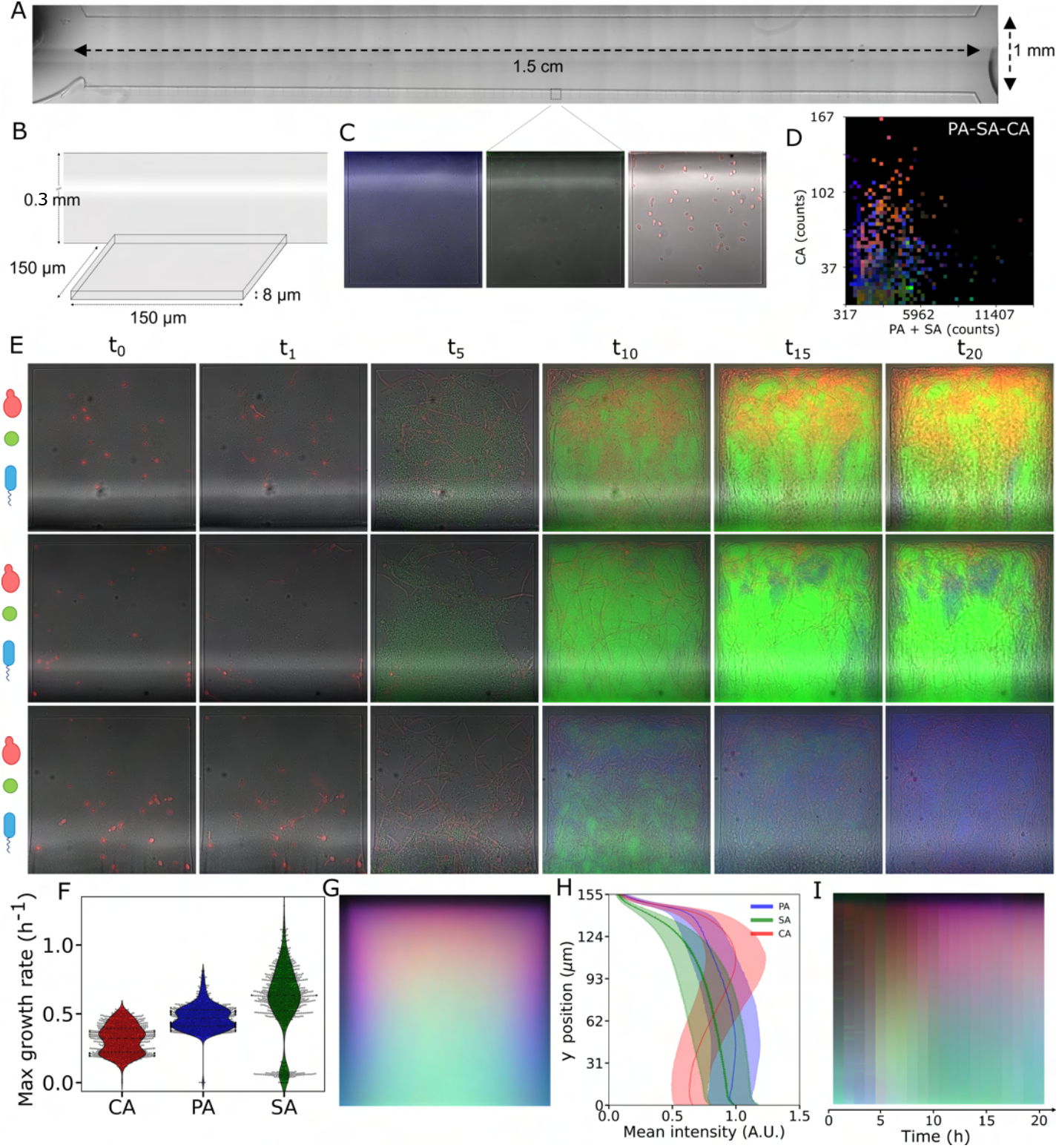
Spatial organization and coexistence emerge in a channel that mimics the distal lung. a) architecture and dimensions of the microfluidic chip. b) zoom-in to the dimensions of the quasi 2D alveoli-like spaces. c) *P. aeruginosa* (left, blue), *S. aureus* (center, green), and *C. albicans* (right, red) at the seeding. They express CFP, GFP and dTomato, respectively. d) mean fluorescence in the distal 25% of the microchambers (colour) after 20 hours, plotted as a function of the initial seeding conditions. e) example time traces from different experiments of the PA-SA-CA community growing in the alveoli-like spaces. f) maximum growth rates extracted from the time course of the different fluorescence channels (9 experiments, 900 boxes). g) average behavior at 20 hours, obtained by summing up all of the boxes (9 experiments, 900 boxes). h) fluorescence intensity profiles of the three fluorescence channels, proxy for PA, SA and CA abundance, along the vertical axis. i) kymograph showing the progression of the average behavior over 20 hours of observation. Each band is obtained by averaging the mean image (g) along the x axis.

**Figure 3.**
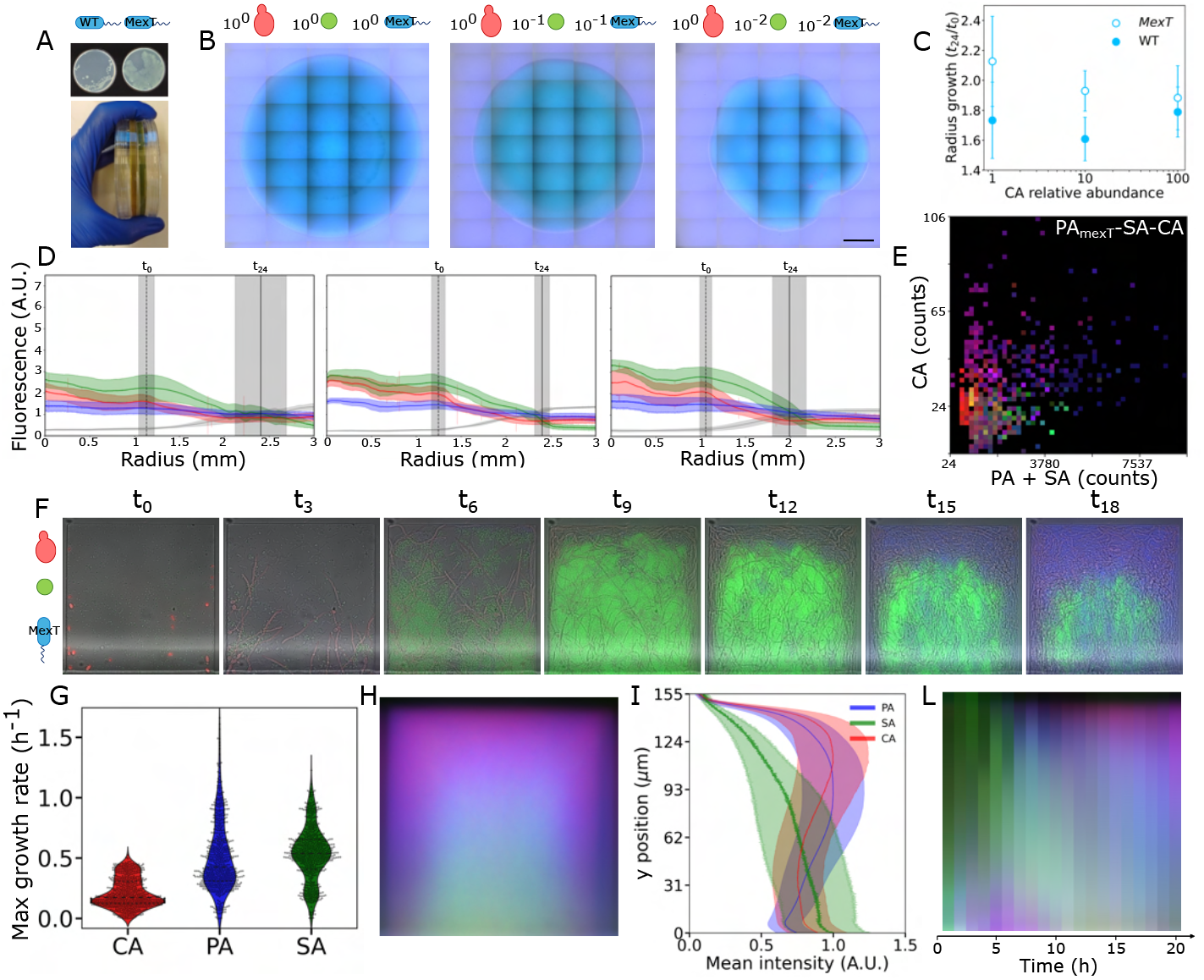
An hypervirulent strain of *P. aeruginosa* with a mutated *mexT* does not significantly alter spatial organization. a) the *mexT* mutant shows a characteristically enlarged colony size and secretes large quantities of piocyanin. b) PA_*mexT*_-SA-CA polymicrobial community seeded at various ratios (left 1:1:1, center 0.1,0.1,1, right 0.01,0.01,1). The *P. aeruginosa* strain expresses the mutated *mexT*. Scale bar = 1 mm. c) comparison of radius growth between PA-SA-CA communities with a reference strain of *P. aeruginosa* (filled circles) or the *mexT* mutant (empty circles). The error bar shows the standard deviation between repeats (6 colonies for a PA-SA-CA initial titer of 1:1:1, 3 colonies for 0.1,0.1,1, and 6 colonies for 0.01,0.01,1. d) average intensity profiles for red, green, blue (fluorescence) and gray (brightfield) channels. Gray indicates brightfield intensity as a proxy for total colony size. Traces are obtained as the average of 360 radii, one every angular degree of the colony. The shaded areas show the standard deviation. The vertical black lines indicate the average radius of the colony at the start and after 24 hours. The shaded area shows the standard deviation. (left) PA-SA-CA ratios 1:1:1, center (0.1,0.1,1), right (0.01,0.01,1). e) mean fluorescence in the distal 25% of the microchambers (colour) after 20 hours, plotted as a function of the initial seeding conditions. f) example time trace of the 3-species polymicrobial community containing the *mexT* mutant in the alveoli mimics. g) growth rates extracted from the timelines of the fluorescence channels (6 experiments, 600 boxes). h) average behavior at 20 hours, obtained by summing up all of the boxes (6 experiments, 600 boxes). i) fluorescence intensity profiles of the three fluorescence channels, proxy for PA, SA and CA abundance. l) kymograph showing the progression of the average behavior over 20 hours of observation. Each band is obtained by averaging the mean image (h) along the x axis.

**Figure 4.**
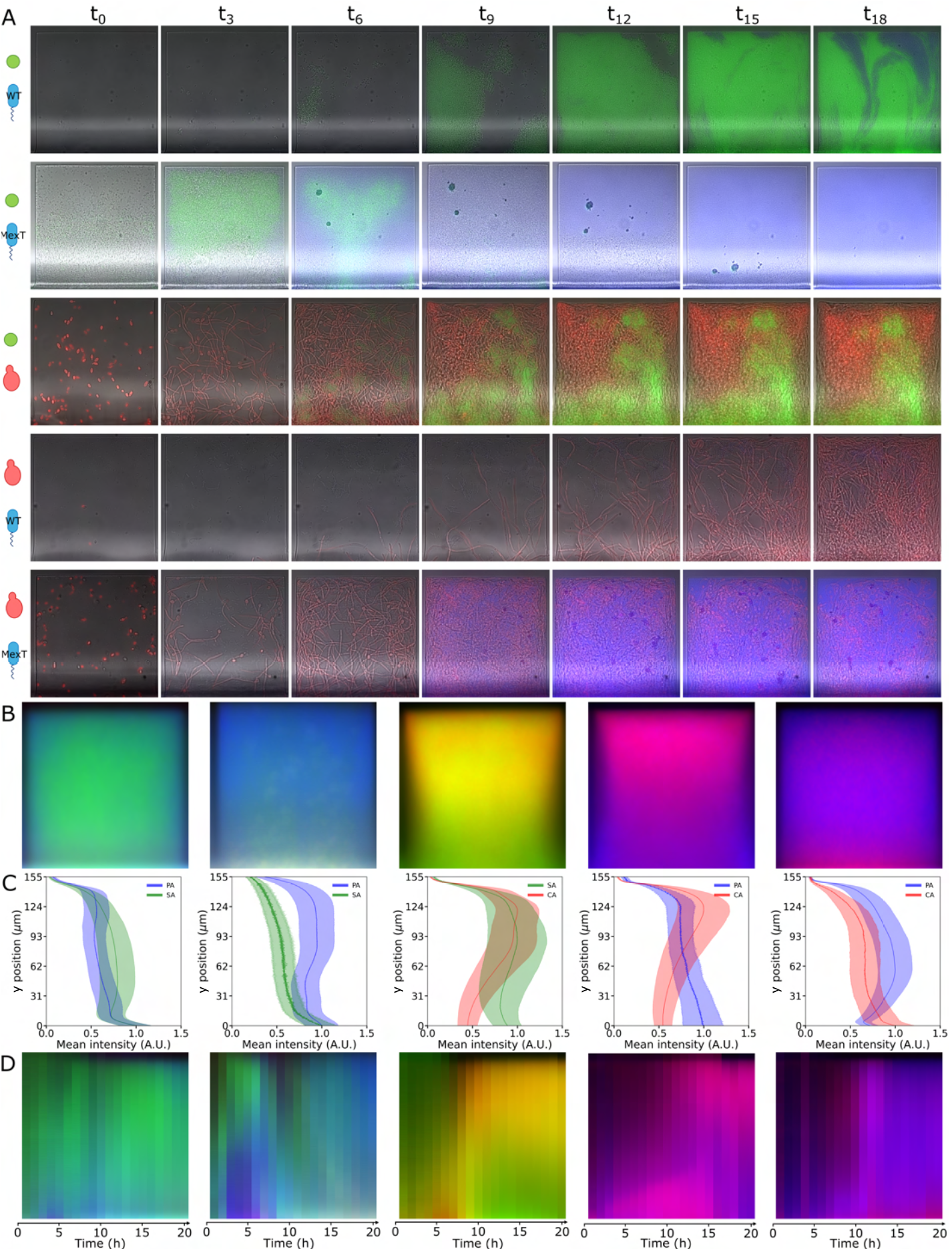
*C. albicans* is necessary for spatial organization. a) example time series of PA-SA, PA_*mexT*_ - SA, SA-CA, PA-CA and PA_*mexT*_ -CA communities. b) average behavior at 20 hours, obtained by summing up all of the boxes (3 experiments, 300 boxes each). Spatial organization emerges also in the absence of SA or PA, but not if PA has a mutated *mexT*.c) fluorescence intensity profiles of the three fluorescence channels, proxy for PA, SA and CA abundance. d) kymograph showing the progression of the average behavior over 20 hours of observation. Each band is obtained by averaging the mean image (b) along the x axis.

**Figure 5.**
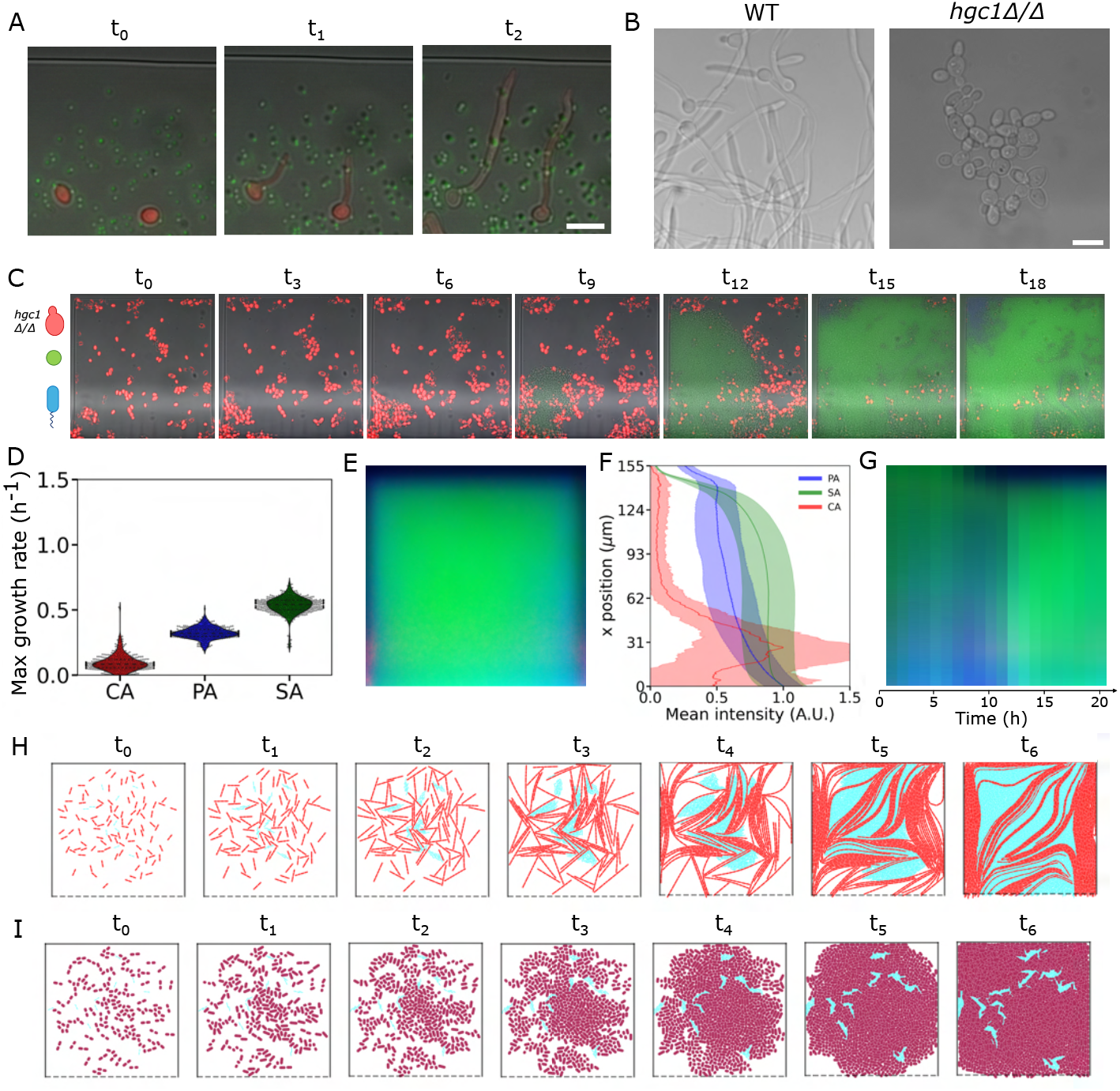
Hyphal growth is necessary for the emergence of spatial organization and coexistence. a) Representative transition from yeast to hyphal form of *C. albicans* within alveoli mimics. b) The yeast-locked mutant *hgc1*Δ:Δ does not grow hyphae in our conditions. c) example time lapse of the *hgc1*Δ:Δ growing in the presence of *P. aeruginosa* and *S. aureus*. C. albicans is segmented from brightfield and false coloured as explained in the methods section. d) growth rate of the 3 species grown in the alveoli mimics. e) average behavior at 20 hours, obtained by summing up all of the boxes (5 experiments, 500 boxes each). f) fluorescence intensity profiles of the three fluorescence channels, proxy for PA, SA and CA abundance. g) kymograph showing the progression of the average behavior over 20 hours of observation. Each band is obtained by averaging the mean image (e) along the x axis. h) 2D agent simulation showing that hyphae-competent *C. albicans* positions itself at the edges. i) 2D agent simulation showing that the yeast-locked mutant *hgc1*Δ:Δ does not enable spatial organization, even when it grows at the same growth rate of the reference strain. Error bars = 10 *µ*m.

### Spatial organization is robust to changes in the virulence of members of the community

To test the robustness of the emerging spatial organization in the community against changes in the virulence of members of the community, we replace the *P. aeruginosa* PA01 reference strain with a *mexT* mutant isolated from the same background. Inactivation of *mexT* is a common feature in clinical isolates, leading to increased virulence (75). The *mexT* mutant tends to form large rough colonies, which express significant amounts of a green pigment called pyocyanin (76) (Fig. 3A). Microscopy of the colonies on ASM agarose surfaces shows increased fitness of the mutant compared to the already dominant reference strain (Fig. 3B, 3C and 3D). *mexT* polymicrobial colonies grow larger (Fig. 3C), with *C. albicans* outcompeted even when 100 times more abundant than *P. aeruginosa* and *S. aureus* at the seeding (Fig. 3B and 3D). In the alveoli-mimics with the *mexT* mutant, seeding communities at different species titers (Fig. 3E), again, shows that spatial organization emerges robustly across 6 independent experiments and 600 alveoli (Fig. 3F and 3H). In this case, the exclusion of *P. aeruginosa* from the closed end is less severe, but more severe for *S. aureus* (Fig. 3I). Although the mutant has a slightly higher growth rate than the reference strain (Fig. 3G), spatial organization still emerges within 10 hours (Fig. 3L) and at the same seeding titers (Fig. 3E).

### Species composition influences spatial organization

Having characterized the behavior of a 3-species community, we investigate the contributions of single members by studying pair interactions. When grown in the absence of *C. albicans, S. aureus* and the *P. aeruginosa* reference strain tend to share the niche (Fig. 4). The *mexT* mutation alters this balance, resulting in a characteristic succession pattern in which *S. aureus* initially grows robustly to then be removed from the box by *P. aeruginosa* that systematically conquers the edges (Fig. 4A, second row). In the presence of *C. albicans*, the absence of either *S. aureus* or *P. aeruginosa* shows little impact on spatial organization, with the yeast preferentially occupying the edges and the closed end of the alveoli mimics as in the three-species scenario (Fig. 4). Similar to the *P. aeruginosa*-*S. aureus* pair, *mexT* shifts the balance towards *P. aeruginosa* (Fig. 4).

### *C. albicans* conquers the closed end of confinements by growing eccentrically with hyphae

These results show that *C. albicans* tends to occupy the edges of confinements and the closed end of our alveoli mimics. This could be the main driver of spatial organization. We therefore seek to investigate how the slowest growing species can have such a dominant role in the ecology of a pathogenic polymicrobial community. While *C. albicans*’s biomass grows overall more slowly in the alveoli mimics, its transition to hyphae in ASM focuses all growth in highly eccentric directions by virtue of the long asymmetric morphology of these structures (Fig. 5A). While growth on a surface is known to stimulate hyphal transition, the ASM used here possesses all the necessary and sufficient conditions to induce hyphal morphogenesis (SI Fig. 6). In confinement, this allows it to rapidly reach the edges of the alveoli-mimic walls and align along them. To test whether hyphal growth is necessary for spatial organization, we repeated the polymicrobial experiments using a yeast-locked *C. albicans* strain that lacks the HGC1 gene, that encodes a cyclin-related protein necessary for hyphal transition in ASM (Fig. 5B). This abolishes the spatial organization observed with the hyphae-competent strain (Fig. 5C, 5D, 5F and 5G). However, the *hgc1*Δ:Δ strain has a slower growth rate than the reference strain (SI Fig. 5). To further test whether the highly eccentric growth conferred to *C. albicans* by its hyphae determines spatial organization, we generate a mutant with an identical growth rate *in silico*, using 2D agent-based simulations. A 2D alveoli mimic is populated by two species with growth rates and cell sizes matching those observed in experiments for the reference strain *C. albicans* and bacteria (grouping together *P. aeruginosa* and *S. aureus*, see *Methods* section). Strikingly, the hyphal *C. albicans* reside at the edges of the alveoli mimic, reproducing the spatial organization observed in experiments (Fig. 5H) We switch the reference *C. albicans* with a *hgc1*Δ:Δ-like strain that does not form elongated cells but has the same growth rate. In this second scenario, the species mix without producing a specific organization (Fig. 5I). These results demonstrate that spatial structure within the alveoli mimics is driven by the mechanical asymmetry of *C. albicans* growing hyphae, which allow them to extend to the walls and corners of the mimic.

## DISCUSSION

By comparing agarose surfaces and confining microfluidic alveoli mimics, we have shown that spatially structured niches can drastically alter ecological outcomes in pathogenic microbes. The 2D alveoli mimics reveal that, within 10 hours after seeding, a polymicrobial community of *P. aeruginosa, S. aureus* and *C. albicans* acquires and maintains a characteristic spatial organization that is not achieved on unconfined planar surfaces. The pivotal factor in this behavior is the capability of *C. albicans* to line the edges of the confined space with its hyphae, excluding the other members of the community. This strategy allows the fungus to survive in the niche despite its slow growth rate, coexisting with the faster growing *S. aureus* and *P. aeruginosa*. In contrast, in the absence of spatial limits on a planar surface, *P. aeruginosa* quickly outcompetes *C. albicans* - even when starting from a hundredfold numerical disadvantage. Like in real alveoli, the edges of the alveoli mimics mediate the contacts between microbes and substrate. In real alveoli, this contact is negotiated by the respiratory epithelium. Therefore, due to its tendency to line confined spaces, *C. albicans* may mediate most of the physical contact between the host and the polymicrobial community *in vivo*. This could have repercussions on treatment design, for example because targeting *C. albicans* might lead to its replacement with more pathogenic microbes. Instead, strategies that blunt *C. albicans*’ toxicity without removing it from the niche may protect the host from further damage.

The robustness of these observations was tested with a *P. aeruginosa mexT* mutant, an adaptation that is frequently found in clinical isolates (75). *mexT* is a transcription regulator that controls the expression of more than 40 genes (77). Its deletion increases virulence by upregulating motility and the production of secreted virulence factors, such as pyocyanin (76). Although in the three-species experiments considered here, the mutant does not qualitatively alter the overall spatial organization trend, it does show increased ecological competitiveness. This is evident from a comparison between Fig. 2G and Fig. 3H with the mutant tending to a distribution that is skewed towards the edges of the microenvironments. Its increased ecological competitiveness manifests fully in the two-species infections, where the *mexT* mutatnt is able to completely eradicate *S. aureus* and limit *C. albicans*’ number. Because its growth rate is similar to that of the reference strain (SI Fig. 5), and those of *S. aureus* and *C. albicans* are not particularly affected by its presence, we hypothesize that its advantage stems from increased motility. This could allow the bacterium to make its way around *C. albicans*’ biomass and dislodge *S. aureus* from crevices, pushing it away through growth.

Taken together, our findings demonstrate that traits controlled at the level of phenotype, such as *C. albicans*’ morphological changes and *P. aeruginosa*’s virulence, can have a significant impact on ecological outcomes when observed in realistic settings to the point of outweighing growth rate alone. Because such evolutionary adaptations are prevalent among microbes and microscale spatial structure is widespread across different environments, we expect these findings to be generalizable to a range of other contexts.

## METHODS

### Microbial strains, media and preculture conditions

For *P. aeruginosa*, we used two strains: a PA01 reference strain expressing CFP (78) from the chromosome and a spontaneous *mexT* frameshift mutant emerging in the same background (and therefore also expressing CFP). We characterize the strains using whole genome sequencing (MicrobesNG, UK). For *S. aureus* we use a SH1000 strain expressing GFP from the chromosome (79). Our reference strain for *C. albicans* is CAF2.1-dTom-NATr that expresses dTomato under the control of the pENO1 enolase promoter (80). The yeast-locked strain is the WYZ12.2 *hgc1*Δ*/*Δ, a CAF2.1 derivative from (81). Prior to any of the experiments, single cultures of *P. aeruginosa, S. aureus* and *C. albicans* are grown overnight in Luria-Bertani broth (0.5% yeast extract, 1% Bacto Tryptone, 0.05% NaCl) in a shaking (220 rpm) incubator at 37°C. 1 ml aliquots of each culture were washed twice in PBS (0.8% NaCl, 0.02% KCl, 1.44% Na2HPO4, 0.24% KH2PO4, pH 7.4) with centrifugation (8000xG, 1 min). The single colonies used to inoculate the overnights are streaked on LB plates from frozen stocks on the day before inoculation for *P. aeruginosa* and up to a week before for *S. aureus* and *C. albicans*. Except where otherwise indicated, all of our experiments are carried out in ASM (saturating amount (*<* 5 g/L) of mucin from porcine stomach type-II dissolved in PBS, saturating amount (*<* 4 g/L) of fish sperm DNA, 1.3 mM NaH2PO4, 1.25 mM Na2HPO4, 0.348 mM KNO3, 0.271 mM K2SO4, 2.28 mM NH4Cl, 14.94 mM KCl, 51.85 mM NaCl, 10 mM MOPS, 1.45 mM Serine, 1.55 mM Glutamic acid, 1.66 mM Proline, 1.2 mM Glycine, 1.78 mM Alanine, 1.12 mM Valine, 0.63 mM Methionine, 1.12 mM Isoleucine, 1.61 mM Leucine, 0.68 mM Ornitine, 2.13 mM Lysine, 0.31 mM Arginine, 0.01 mM Tryptophan, 0.83 mM Aspartic acid, 0.8 mM Tyrosine, 1.07 mM Threonine, 0.16 mM Cysteine, 0.53 mM Pheny-lalanine, 0.52 mM Histidine, 3 mM Glucose, 9.3 mM L-lactic acid, 1.75 mM CaCl2, 0.6 mM MgCl2, 0.0036 mM FeSO4, 0.3 mM N-acetylglucosamine, 5 ml Egg yolk emulsion, pH 6.8, filter sterilised). Our ASM is prepared as in (64) and is therefore a modified version of (82; 83; 84).

### Soft agarose experiments

4 mm thin agarose plates are produced by filling the lids of 3.5 cm petri dishes (Greiner, UK) with 3 ml molten 1.5% agarose solutions in ASM. The plates are seeded with 0.2 *µ*l mixtures of the washed *P. aeruginosa, S. aureus* and *C. albicans* at various dilution ratios (Fig. 1 and Fig. 3). To prevent agarose marking with the tip of the pipette during seeding, we slowly pipette out the small culture volume until a drop is visible at the end of the tip. The edges of the plates are secured to a custom-made microscope mount with double-sided tape and immediately imaged using a Nikon 10× Plan Apo air objective with a numeric aperture of 0.45 on a Nikon Ti-E microscope using Nikon Perfect Focus System. A total of 280 images per colony were collected using a Teledyne FLIR BFS-U3-70S7M-C with a 7.1 MP Sony IMX428 monochrome image sensor. All images are captured at 3208×2200 pixels, with a resolution of 0.43 *µ*m/pixel. Each FOV is imaged 7 times, of which 1 in brightfield and 3 in 3 fluorescence channels, each imaged at two different exposure time and gain combinations: 0.5 s, 20 gain and 0.2 s, 10 gain for dTomato; 0.2 s, 32 gain and 0.005 s, 32 gain for GFP; 0.06 s, 32 gain and 0.005 s and 32 gain for CFP. Illumination is provided by color Light Emitting Diodes: red for brightfield red (Lumileds Luxeon Z LXZ1-PD01), indigo for CFP indigo (Lumileds Luxeon UV LHUV-0425), blue for GFP (Lumileds Luxeon Z LXZ1-PB01), and lime for dTomato (Lumileds Luxeon Z LXZ1-PX01). For fluorescence imaging we use Semrock optical filter sets (IDEX Health and Science, USA), CFP-2432C for CFP, GFP-3035D for GFP and TxRed-4040C for dTomato. After imaging, which typically took 2 hours, positions of the colonies on the plates are recorded and the plates are temporarily sealed with a second lid glued with Covergrip coverslip sealant (Biotium, USA) to maintain moisture. The plates are transferred to a static 37°C incubator for 24 hours and imaged again.

### Microfluidic lung mimic production

Microfluidic master molds are produced in-house using SU8 (Kayaku AM, USA) soft-lithography following the manufacturer’s guidelines. A first thin layer of SU8-2010 is deposited on a 10.16 cm (4 inches) silicon wafer (PI-KEM, UK), exposed to UV light in a MicroWriter ML3 Pro (Durham Magneto Optics, UK) and baked to obtain a final thickness of approximately 8 *µ*m (Fig. 2C). A second layer of SU8-2075 is then deposited on the mold, exposed to UV light, baked and washed with SU8 developer (Kayaku AM, USA) to obtain the main trench (Fig. 2A), with a height of 0.3 mm. The designs are inspired by the family machine (74). The molds are then used to fabricate microfluidic chips using degassed PDMS at a 1:10 ratio between curing agent and elastomer. Curing of the PDMS was performed in a 60°C oven. Before use, chips are released from the mold using a scalpel, inlets and outlets are punched using a biopsy puncher with 0.77 mm internal diameter (World Precision Instruments-Europe, UK), the chip is plasma bonded to a 0.145 mm thick glass coverslip (VWR, UK) and baked at 60°C for at least 10 minutes. For the microfabrication of a mother machine that could house *C. albicans*, we use the same procedure, but lower the height of the first SU8 layer to 5 *µ*m.

### Lung-mimic experiments preparation

Microfluidic channels are loaded with microbes mixtures obtained from the washed overnight cultures and inspected using microscopy. To carry out the experiment, inlets and outlets of the microfluidic chips are connected to Tygon tubing (Cole Parmer; 0.020” × 0.060” outside diameter), using 90° bent connectors (Intertronics, UK; stainless steel, 0.89-0.58 OD-ID, 20 gauge) as before (85). A 20 *µ*/min flow of ASM medium is prompted by a syringe pump (kdScientific, USA), loaded with a 60 ml plastic syringe (BD plastipak, UK), connected to the tubing via blunt needles (Intertronics, UK; stainless steel, straight blunt, 1/2”, 23 gauge).

### Lung-mimic experiments

Experiments are carried out at 37°C by housing the microfluidic chip in custom-built heating elements that are fitted to the automated xy stage of a Nikon Eclipse Ti-E. Before starting the experiment, and after loading, an arbitrarily fast flow pulse of fresh ASM medium is delivered to the main trench of the chip to remove microbes that had not entered in the alveoli-mimicking boxes. Imaging is automatically carried out in two phases. In the first phase, we imaged the 100 boxes on 1 side of the channel using a Nikon 40× CFI Plan Fluor air objective with a numeric aperture of 0.75. Each box is imaged 19 times, once in brightfield and 18 times in fluorescence using the same LEDs and filters given in the *Soft agarose experiments* section. Each fluorescence channel is imaged at maximum gain at three different z offsets, with two exposure times: 0.4 s and 0.1 s for dTomato; 0.05 s and 0.003 for GFP; 0.5s and 0.05s for CFP. In the second phase, we image the whole channel using the 10x air objective described in the *Soft agarose experiments* section. At this stage, we collect 266 images per channel for a total of 38 FOVs, each imaged 7 times: once in brightfield and twice per each fluorescence channel, using 32 gain and different exposure times: 0.5 s and 0.2 s for dTomato; 0.05 s and 0.005 s for GFP; 0.06 s and 0.005 s for CFP. The two phases combined typically take 54 minutes of microscope time and are automatically repeated every hour.

### Mother machine experiments

For the mother machine experiments to evaluate the correspondence between biomass and fluorescence, we use two mother machines: one with pistons with width and height below 2 *µ*m (85) for *P. aeruginosa* and *S. aureus* and one with pistons with width and height below 6 *µ*m for *C. albicans*. Before loading the cells in the pistons, the channels are incubated overnight at 37°C with a 2% solution of bovine serum albumin (BSA). Loading is carried out from 50x concentrated LB overnight cultures, double-washed in PBS in all cases. For *P. aeruginosa* and *S. aureus* loading occurred spontaneously, while for *C. albicans* we enhance it by spinning the mother machine at 1000 rpm in a spin coater for 1 minute. Once loading is satisfactory, the inlet and outlet of the mother machine are attached to a syringe pump, as described above, using a 5 ml syringe (BD plastipak, UK). The chip is heated to 37°C and ASM is flowed in at 5 *µ*l/min.

### Simulations

Simulations utilize an agent-based model based on (86), in which cells grow in a two-dimensional plane. Bacterial cells are designated as a single species to represent both PA and SA; symbolically, species *S* = PA-SA. *Candida* hyphae are represented by chains of linked cells, of species *S* = CA (SI Fig. 7).

Cells grow linearly from an initial length *ℓ*_*S*_ with an average growth rate *µ*_*S*_, while cell diameters *d*_*S*_ are constant. The instantaneous length of each cell *i* is *ℓ*_*i*_(Δ*t*) = *ℓ*_*S*_ + *µ*_*i*_Δ*t*_*i*_ at a time Δ*t*_*i*_ since division for an individual growth rate *µ*_*i*_ drawn uniformly from (*µ*_*S*_*/*2, 3*µ*_*S*_*/*2) at division. Cells grow until they reach a set division length 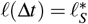,at which point they divide into two daughters of equal length. Daughter cells inherit their mother’s orientation with small perturbations.

The position ***r***_*i*_ and orientation ***u***_*i*_ evolve according to overdamped equations of motion

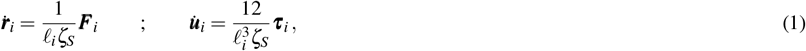

where *ζ*_*S*_ is the friction per unit length, and ***F***_*i*_ and ***τ***_*i*_ are the net force and torque on the *i*^th^ cell (87; 86). Cells are simulated as spherocylinders that interact through steric pair potentials and are subject to 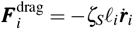 and 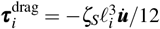,which approximates drag as orientation-independent and proportional to cell length (88). All other forces and torques result from pair potentials *U*_*i j*_ as 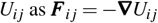. Steric potentials between spherocylinders *i* and *j* are modelled via a purely repulsive Lennard-Jones potential (89)

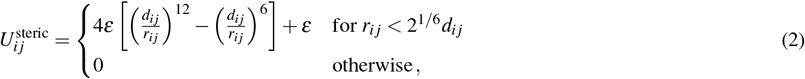

with repulsion strength *ε*, characteristic size *d*_*i j*_ set to the mean diameter of the pair of cells and the separation *r*_*i j*_ calculated between two fictitious spheres located at the points of closest approach along the axes of symmetry of the two spherocylinders.The torque on cell *i* due to contact with *j* is 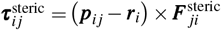, where the location of the force ***p***_*i j*_ is taken to be at the surface of cell *i* closest to the axis of cell *j*.

In addition, each *C. albicans* cell has bonds that connect it to the extreme ends of the axes of adjacent cells in the hyphal chain. The bond length *l*_ij_ obeys a potential 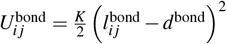 for a compression modulus *K* and rest length *d*^bond^, which is set to be slightly smaller than *d*_*S*_ for *Candida* to avoid the smaller bacterial cells penetrating the chain.

The bending energy is 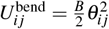 for bending stiffness *B*, with *θ*_*i j*_ the angle between the axes of neighboring segments on the chain. The torques result from applying the forces at the ends of the cell axes. Hyphae filaments divide when the number of segments reaches a maximum value of *N* = 50; at which point, the middle bond is removed.

The bacterial parameters (*S* = PA-SA) are 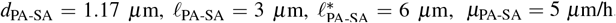,*ζ*_PA-SA_ = 200 Pa h. The parameters for *C. albicans* (*S* = CA) are 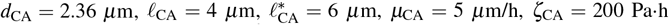. The hyphae compression modulus is *K* = 4 *×*10^*−*6^ N/m, bending stiffness *B* = 4*×* 10^*−*17^ N·m and the rest length *d*^bond^ = 0.65 *d*_*CA*_. For both bacteria and *C. albicans*, the steric repulsion is set to *ε* = 5.5 *×*10^*−*12^ N·m. A single microfluidic alveoli mimic is simulated as a 2D square microchamber of length *L* = 150*µ*m. The microchamber is constructed of three planar walls and a planar opening. The walls are modeled as repulsive potentials that increase quadratically with the degree of cell/segment overlap with modulus *K*^wall^ = 4 *×* 10^6^ N/m. When both ends of a cell pass the opening plane, it is instantaneously removed from the simulation. This simulates the loss of cells to flow in the main microfluidic channel.

### Data analysis

The code used for image analysis and statistics is available at: 10.5281/zenodo.15005092.

### Image correction and preprocessing

Image processing is performed in Python, Fiji (90) and ilastik (91). Brightfield and fluorescence images from ASM-agarose surfaces (Fig. 1) are normalized by the illumination profiles of the respective light sources before stitching. The profiles are obtained by averaging the signal from at least 15 FOVs captured on a coverslip without any objects in focus. In all cases, stitching is performed using a custom-made Python script with the exception of Fig. 2A, for which we use the Grid/Collection stitching plugin by (92) in Fiji. Because the fluorescence change is relatively small at the center of the images where the alveoli-mimicking boxes are captured, we choose not to perform such illumination profile correction in any of the other microscopy images presented in the text. The FOV of the camera with the 40x air objective is 382×262 *µ*m and therefore we are able to image two boxes at a time. To extract single boxes, we use a Python-enhanced manual approach in which, for each image, frames of 1400×1400 pixels are superimposed to the brightfield images and their position manually fine-tuned to match the microfluidic boxes. This is done for the 100 positions of the boxes. Choosing a frame size that is slightly larger than the box, coupled with the robustness of the stage control allows us to bypass image registration. During experiment setup, we also strived to minimize tilt along the channel, which allows us to neglect rotational adjustments at the time of analysis. RGB images of single FOVs are generated in Fiji by merging the original 16-bit images in the gray, red, green, and blue channels. We take fluorescence images at three offsets in the box: top and bottom surfaces and middle. For the presentation of single FOVs, we use the bottom offset (glass surface) as this captured most of the *P. aeruginosa* and *S. aureus* cells in the early time points and the central one for the fluorescence channel of *C. albicans*. The final images are exported as 24-bit RGB-color images. The average images (Fig. 2F, 3F, 4B, 5E) are obtained by adding together in the RGB channels the fluorescence images of 100 boxes per experiment per time point. To capture the total behavior in the box, we use the central offsets for all channels. The values are normalized by the maximum value found for the specific fluorescence channel across the entire time series. In our large dataset (*>*1.5M tiff images across 50 experiments for the microfluidic experiments alone), we capture rare events in which the microscope-camera combination fails to acquire single frames. When an image at a given offset is found missing or corrupted, we replace it with the closest offset at the same position. When none of the alternative offsets are available, we replace the image with its closest available preceding time point. Kymographs are obtained by averaging the sums of 20 consecutive time points along the image’s x-axis. To enhance visibility and produce square kymographs, each time point is presented as a 70-pixel wide column. Before plotting the fluorescence profiles (Fig. 2G, 3H, 4C and 5F) along the y-axis of the summed images, we apply the illumination profile correction explained above. Because the *hgc1*Δ:Δ *C. albicans* strain shows a reduced growth rate, we choose not to burden it further with the expression of dTomato. Instead, segmentation is carried out on the brightfield images using ilastik (91). Pixels belonging to the cells (foreground) and the background are assigned arbitrary fluorescence values of 20000 and 1000, respectively, matching those measured in the fluorescent reference strain.

### Data extraction from colonies

Colonies are manually segmented in Fiji (90), with all of the subsequent steps carried out in Python. To estimate the mean colony radius, we calculate the distance of each point on the perimeter from the centroid using 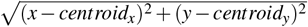. The centroid positions are estimated using OpenCV (93). To extract fluorescence and brightfield values, we split the colonies in 360 sections, each 1° large. For each section, a radius is drawn and gray values are extracted along the radius for each of the four imaging channels (1 brightfield and 3 fluorescence channels).

### Growth rate extraction

To extract growth rates from fluorescence time courses, after having observed that fluorescence reflects biomass well (SI Fig. 4), we compute the mean fluorescence values from illumination corrected boxes. Each of the resulting growth curves is independently fit using (94) extracting the maximum growth rate, with the following ranges of fitting parameters: amplitude -5,5; flexibility -6,2; error -5,2.

### Biomass to fluorescence relationship

To test whether fluorescence is a good proxy for biomass, we extracte fluorescence values from mother machine pistons. We examine 13 FOVs or more for *P. aeruginosa* and *S. aureus*, with around 70 pistons each and 42 for *C. albicans* with 35 pistons each. We did not perform single cell segmentation. Instead, fluorescence images are segmented using the adaptiveThreshold function with the mean thresholding method (block size = 71, constant = 4) of OpenCV (93). We then apply global thresholding using Otsu’s method. Segmented images are the result of the intersection of the two methods, to which 3 rounds of median blurs are applied. Fluorescence per time is calculated as the total mean of the fluorescence signal per pixel.

### Single cell segmentation

The initial titers of cells in the alveoli mimics are extracted using a custom-made segmentation algorithm in Python. Using OpenCV (93), images are thresholded using the adaptiveThreshold function with the mean thresholding method. Contours are extracted from the resulting binary images and filtered for size to eliminate noise and debris. To separate large cell masses, the solidity of the contours is evaluated by estimating their hull size and split recursively through the center when a certain size limit is exceeded. This is sufficient to differentiate single *C. albicans* cells, but for *S. aureus* and *P. aeruginosa* we add a further splitting step that counts the number of cells within larger, still unsplit blobs. For *C. albicans*, before finding the final contours and quantifying their characteristics (position and sizes), we perform a dilation step to fill small gaps. The *P. aeruginosa* strain used produces the dimmest fluorescence signal per cell and thus its analysis requires some additional steps. Each image is copied twice, and the resulting three images (A, B and C) undergo slightly different processing steps: “A” is thresholded as before; “B” is smoothed using the Gaussianblur method before thresholding, and “C” is contrast-enhanced using Contrast Limited Adaptive Histogram Equalization. The foregrounds of the three images are combined and further processed for contour individuation as done for *S. aureus*. We test the efficacy of this approach by benchmarking the numbers of cells segmented against counts obtained through visual inspection obtaining F1-scores equal or above 95% (SI Table 1).

## AUTHOR CONTRIBUTIONS

L.M. Conceptualization, Investigation, Funding acquisition, Data curation, Formal analysis, Investigation, Methodology, Software, Validation, Project administration, Writing – original draft, Writing – review and editing. L.S. Investigation, Software, Data curation, Formal analysis, Writing – review and editing. R.C. Software, Writing – review and editing. J.K. Methodology, Resources, Writing – review and editing. E.B. Resources, Writing – review and editing. C.M. Resources, Writing – review and editing. T.N.S. Supervision, Funding acquisition, Software, Writing – review and editing. A.B. Supervision, Funding acquisition, Software, Writing – review and editing. M.W. Conceptualization, Resources, Writing – review and editing. P.C. Conceptualization, Resources, Writing – review and editing.

## ACKNOWLEDGEMENTS

We thank Bartłomiej Wacław, Teuta Pilizota and Mie Monti for insightful discussions. We are also thankful to Simon Foster, Rob Wheeler and Yue Wang, and Tim Tolker-Nielsen for the gift of the *S. aureus, C. albicans* and *P.aeruginosa* strains, respectively. LM acknowledges funding from the Herchel Smith fund, Postdoctoral fellowship. JK and PC acknowledge funding from the UK CF Trust SRC 016. EB acknoledges funding from the Oliver Gatty Trust. MW was supported by the CF Trust (a Venture and Innovation Award (VIA)) and from the Leverhulme Trust. This research has received funding from the European Research Council (ERC) under the European Union’s Horizon 2020 research and innovation program (Grant agreement No. 851196). For the purpose of open access, the author has applied a Creative Commons Attribution (CC BY) license to any Author Accepted Manuscript version arising from this submission.

## DATA AND MATERIALS AVAILABILITY STATEMENT

All data are available at 10.5281/zenodo.15005092.

