## Supplementary information for "Hyphal growth determines spatial organization and coexistence in a pathogenic polymicrobial community within alveoli-like geometries"

### SUPPLEMENTARY FIGURES

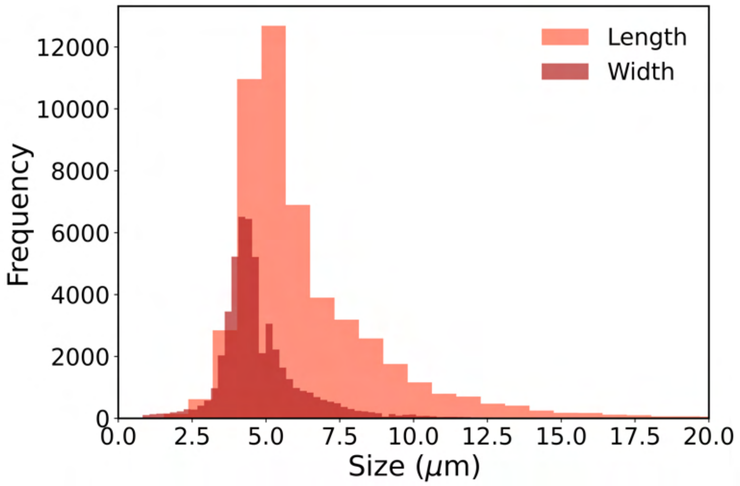


Supplementary figure 1: **Size distribution of *C. albicans* at the seeding.** Measures where extracted via segmentation of the first time point from 9 PA-SA-CA experiments (900 microchambers).

| Species | Automatic/manual count | F1-score |
|---|---|---|
| *P. aeruginosa* | 1145/1042, 1017/1019, 966/1179 | 0.95 |
| *S. aureus* | 1135/1140, 986/917, 743/796 | 0.96 |
| *C. albicans* | 17/17, 4/4, 17/16, 12/11 | 0.98 |

Table 1: **Benchmarking of segmentation algorithms.** The manual count is taken as the true count.

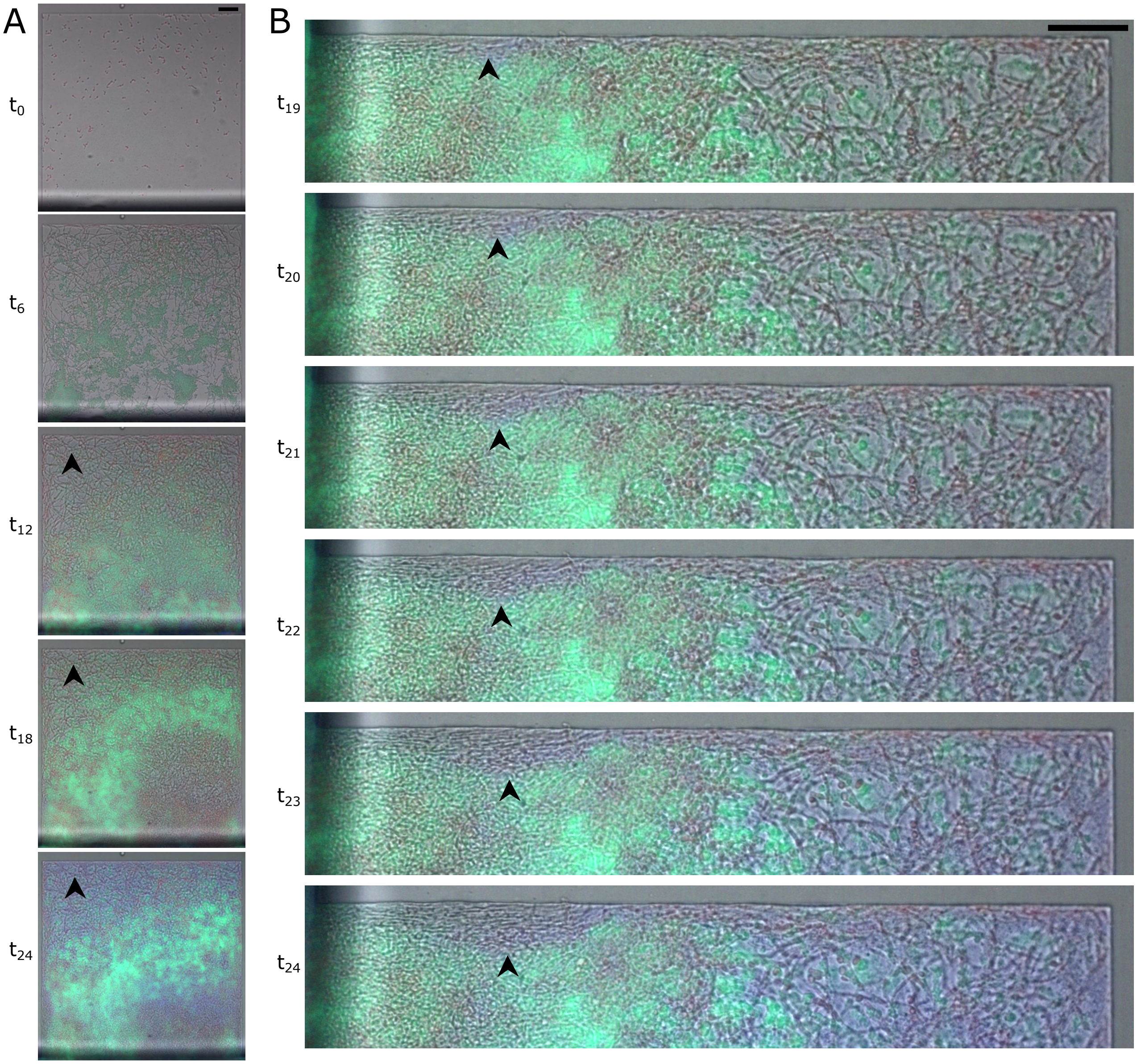


Supplementary figure 2: **Increasing the box volume while maintaining chamber height unchanged leads to lower medium exchange per unit volume and nutritional gradients.** a) Time course of the PA-SA-CA polymicrobial community in a box with side length 500 $\mu$m and height 8 $\mu$m. The *C. albicans* at the closed end of the microenvironment stops growing. The arrowheads point to an area where growth has clearly stopped. b) zoom-in from (a) tilted 90° to the right showing that in the proximity of the opening, *C. albicans* conquers the edge and excludes bacteria. Black arrows indicate the area of progressive growth. Scale bars =50 $\mu$m.

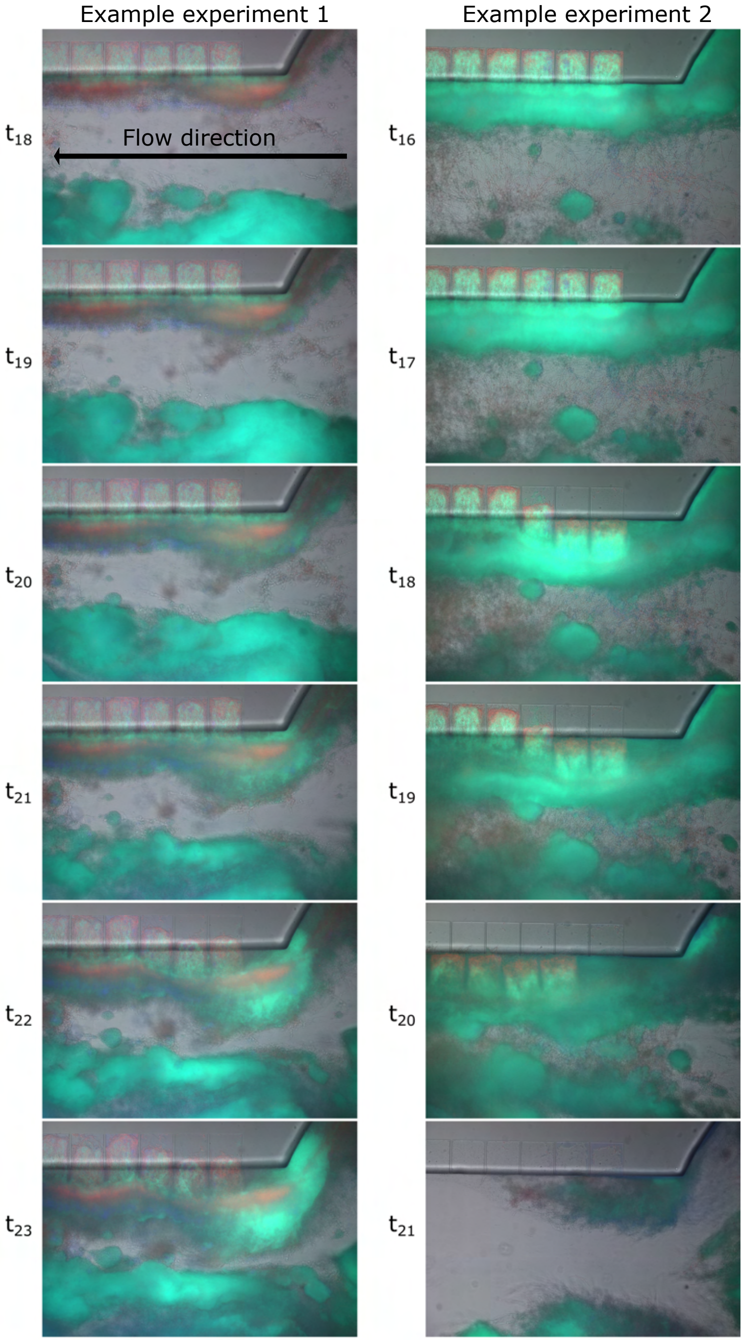


Supplementary figure 3: **Biofilm behavior in the main channel.** After 18 to 20 hours, growth in the main channel becomes substantial and the resulting biomass, further to potentially decreasing perfusion of nutrients in the boxes, can be peeled off by the flow taking away box contents. For this reason we limit our observations to the first 20 hours. Two examples from independent experiments are given (left and right). The size of the boxes' sides is 150 $\mu$m and provides scale.

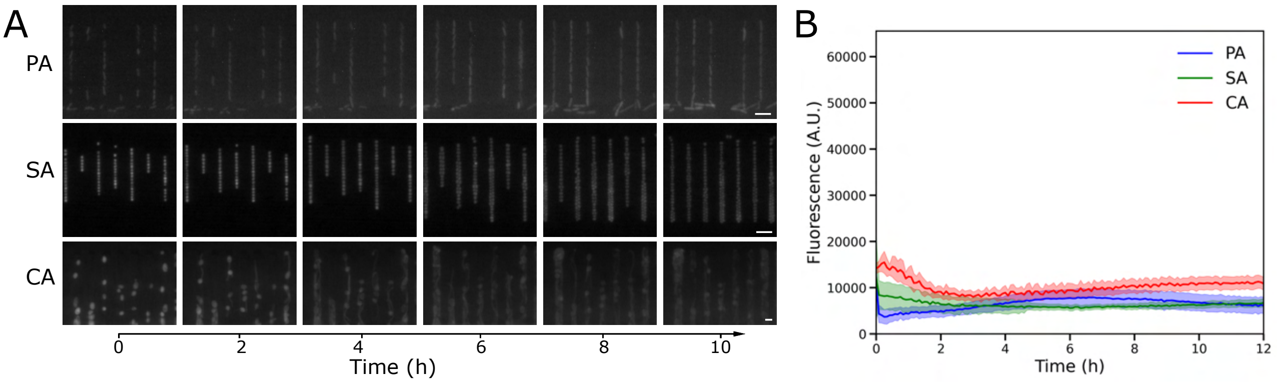


Supplementary figure 4: **Fluorescence is a good proxy for biomass in our strains.** a) fluorescence profiles were extracted from timelapse experiments in mother machines in ASM. b) Average of 13 fovs or more for *P. aeruginosa* and *S. aureus*, with around 70 pistons each and 42 for *C. albicans* with 35 pistons each plotted in the camera sensitivity range. The shaded areas indicate the standard deviation. *P. aeruginosa* = blue, *S. aureus* = green, *C. albicans* = red. Scale bar = 5$\mu$m.

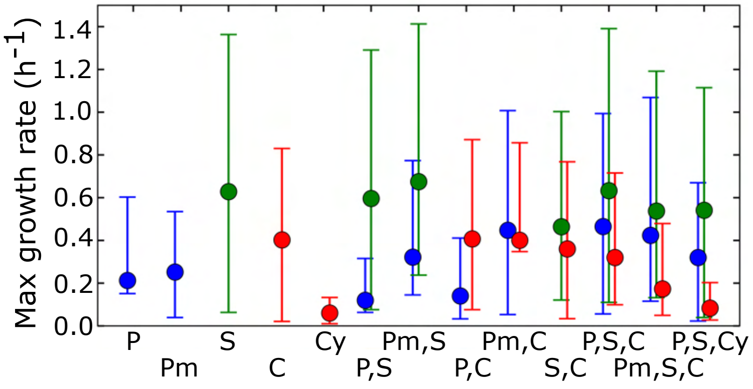


Supplementary figure 5: **Comparisons between growth rates extracted from alveoli mimics.** *Pm* indicates *P. aeruginosa mexT* mutant, *Cy* indicates *C. albicans hgc1*Δ/Δ mutant. *P. aeruginosa* = blue, *S. aureus* = green, *C. albicans* = red. Each average is extracted from at least 300 microenvironments from at least 3 independent experiments, the error bar shows the standard deviation.

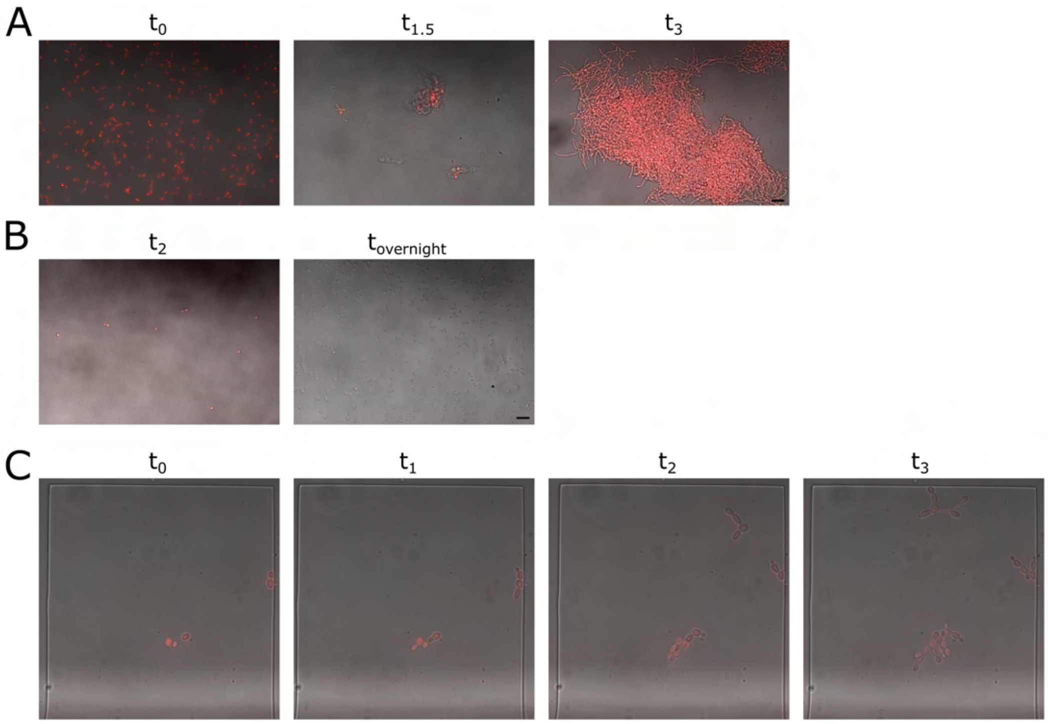


Supplementary figure 6: **Hyphal transition in the microenvironments is not due to the contact with the surface.** a) Time course of *C. albicans* cells grown in shake flasks in ASM at 37°C. In ASM surfaces are not necessary for hyphae formation. Scale bar = 20 $\mu$m. b) *C. albicans* growth in YPD. Scale bar = 20 $\mu$m. c) *C. albicans* growth in YPD in microenvironments and production of pseudohyphae: the microenvironment's surface is not sufficient to induce hyphae formation. The size of the boxes' sides is 150 $\mu$m and provides scale.

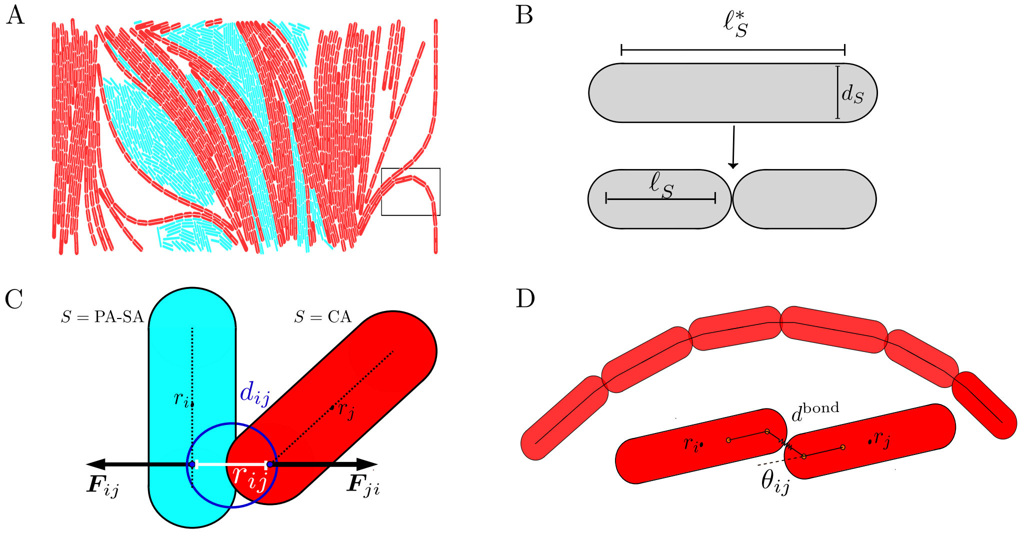


Supplementary figure 7: **Schematic description of the species used in the simulations.** a) Snapshot illustrating bacteria, (S=PA-SA) in cyan, and fungal hyphae (S=CA) in red. b) Schematic for division events of both bacteria cells and hyphae segments. Cells/segments increase their length $\ell$ linearly in time until they reach a critical length, $\ell_S^*$, at which point they divide into two daughters of length $\ell_S$ and diameter $d_S$. c) Steric repulsion forces $\boldsymbol{F}_{ij}$ and $\boldsymbol{F}_{ji}$ between cells/segments $i$ and $j$ results from the nearest distance $r_{ij}$. The characteristic diameter $d_{ij}$ is the average diameter of the interacting segments. D) Magnified view of hyphal segments within the highlighted area (black rectangle in A), showing segments connected by bonds (top). A schematic (bottom) demonstrates two segments chained together. Each CA segment in a chain is connected via bonds linking the extreme ends of adjacent cells in the hyphal chain. The bonds have a rest length $d^{\text{bond}}$ and $\theta_{ij}$ is the angle between the axes of neighbouring segments.
